# Development of noninvasive biomarkers of response to proteasome inhibitor therapy (ixazomib) by imaging disrupted protein homeostasis in mouse models of solid tumors

**DOI:** 10.1101/193623

**Authors:** Yanan Zhu, Rajiv Ramasawmy, Sean Peter Johnson, Valerie Taylor, Alasdair Gibb, R Barbara Pedley, Nibedita Chattopadhyay, Mark F Lythgoe, Xavier Golay, Daniel Bradley, Simon Walker-Samuel

## Abstract

With clinically-approved proteasome inhibitors now a standard of care for multiple myeloma, and increasing interest in their use in solid tumors, methods for monitoring therapeutic response *in vivo* are critically required. Here, we show that tumor protein homeostasis can be noninvasively monitored, using chemical exchange (CEST) magnetic resonance imaging (MRI) as a surrogate marker for proteasome inhibition, alongside diffusion MRI and relaxometry. We show that the *in vivo* CEST signal associated with amides and amines increases in proportion to proteasome inhibitor dose (ixazomib) and the magnitude of therapeutic effect in colorectal cancer xenografts. Moreover, we show that SW1222 and LS174T human colorectal cancer cell lines demonstrate differing sensitivities to ixazomib, which was reflected in our MRI measurements. We also found evidence of a mild stimulation in tumor growth at low ixazomib doses. Our results therefore identify CEST MRI as a promising method for safely and noninvasively monitoring changes in tumor protein homeostasis.

## Introduction

Loss of protein homeostasis is associated with a range of pathological conditions including most forms of dementia (1), amyloidosis (2) and cancers (3). In cancer, malignant cells exhibit an abnormally high turnover of protein (3), which is most notably exploited as a therapeutic target by proteasome inhibitors (PIs) (4). By inhibiting the action of proteasomes, PIs disrupt protein homeostasis, resulting in apoptotic cell death. PIs are now in routine clinical use for treating haematological malignancies (5-7).

Ixazomib (Ninlaro/MLN2238/MLN9708, Takeda Pharmaceuticals Company Limited) is the first orally-available PI therapy (8, 9) with approval for use in the United States, Europe and Japan for combinational treatment in multiple myeloma (9, 10). Ixazomib has also shown promise in solid tumor rodent models, including colorectal tumors, and there is an increasing research interest in this area (8, 11-13).

Here, we propose a new set of noninvasive imaging biomarkers for detecting disrupted protein homeostasis, resulting from PI therapy, based on magnetic resonance imaging (MRI). In particular, we have focused on chemical exchange saturation transfer (CEST) MRI, which has previously been used to detect changes in protein content in cancer (14-16). CEST image contrast can be tuned to reflect the exchange of protons between water and various chemical groups, including amides (e.g. in protein backbones) (17), amines and hydroxyls. This enables the small signal from solutes containing such groups to be amplified via the much larger water pool signal. We hypothesized that successful proteasome inhibition, and associated protein accumulation (4, 18), would result in an increase in CEST signal associated with amide and amine groups.

Alongside CEST, we evaluated a range of quantitative, noninvasive MRI measurements (diffusion MRI, *T*_1_ and *T*_2_ relaxometry), in LS174T and SW1222 xenograft models of human colorectal carcinoma. *In vivo* measurements of acute (up to 72 hours) dosing with ixazomib, at three different dose concentrations, were undertaken and compared with gold-standard histological measures (19-21). Biological validation studies, such as undertaken here, are a vital component of the translation pathway for imaging biomarkers into the clinic (22).

The aims of our study were to: 1) determine the efficacy of ixazomib in colorectal carcinoma cell lines and murine xenografts; and 2) evaluate the ability of noninvasive MRI measurements to evaluate this response *in vivo,* with a view to their translation into the clinic.

## Materials and Methods

Details of cell culturing methods for SW1222 and LS174T cells are provided in the supplemental methods.

### *In vitro* MTT assay

MTT (Vybrant MTT Cell Proliferation Assay Kit, Life Technologies) reagent at 12 mM was prepared with a dilution of 1 mL of sterile PBS to one 5 mg MTT powder, according to manufacturer’s instructions. Before treatment, 10,000 in 100 μL of complete media per well were seeded, for 9 hours, in a clear polystyrene 96-well flat bottom microplate (Greiner Bio-One).

Ixazomib was diluted from 10 mM stock solution in complete media from 400 to 1.6 nM using 2-fold serial dilution. Media in each well was removed and replaced with 90 μL of ixazomib, with corresponding concentration. Cells were left for 24, 48 or 72 hours, then labeled with 10 μL of the 12 mM MTT reagent, and incubated for 2 hours at 37°C. After the incubation period, the media was removed and replaced with 100 μL of DMSO per well, and mixed thoroughly using an orbital plate shaker for 2 minutes. The microplate was incubated for 37°C for 10 min and were read using a microplate absorbance reader at 540 nm.

Cell viability was estimated from the absorbance measure (unitless), a measure of cell viability as a percentage of the mean absorbance of control cells. These data were fitted to the modified Hill equation, from which the *EC*_*50*_ (half-maximal stimulatory concentration) and *IC*_*50*_ (half-maximal inhibitory concentration) were estimated (see supplemental methods for details).

### Animal Models

All *in vivo* experiments were performed in accordance with the UK Home Office Animals Scientific Procedures Act, 1986 and United Kingdom Coordinating Committee on Cancer Research (UKCCCR) guidelines (23). Mice had access to food and water ad libitum. 5×10^6^ SW1222 or LS174T cells were subcutaneously inoculated on the lower right flank of female CD1 nu/nu mice (0.1 mL per injection).

### *In vivo* experimental design

Tumors were allowed to grow for 14-16 days, then randomly assigned to treatment and control groups. The treatment group received a single dose of ixazomib i.v., at 8, 9.5 or 11 mg kg^-1^. The control group received vehicle, consisting of the drug stock solution (2-Hydroxypropyl-β-cyclodextrin, Sigma Aldrich). Treated mice were maintained in a warmed cage using a thermostatic heating pad (Physitemp Instruments, Inc., New Jersey) and an infrared heating lamp (Zoo Med, UK) providing an ambient temperature of 24-26 °C. Hydrogel was provided to prevent dehydration. Tumor volume (*V*) was measured every 2 days using electronic calipers, according to (24) *V* = *w*^2^*l* / 2, where *l* and *w* are the maximal tumor diameter and the diameter orthogonal to this measurement, respectively. The mass of each mouse was also measured using an electric balance (C-MAG HS 7 IKAMAG^®^, Staufen, Germany).

Baseline MRI data were acquired 24 hours before drug administration (0 hours), with follow-up sessions at 48 and 72 hours. Immediately following the final MRI scan, mice were culled via cervical dislocation, tumors were resected and sent for *ex vivo* analysis.

### *In vivo* MRI protocol

MRI data were acquired using a 9.4T scanner (Agilent Santa Clara, CA) with a 39 mm birdcage coil (Rapid MR International, Columbus, OH). Mice were anaesthetized prior to and throughout each scanning session using isoflurane in O_2_ (2.5% for induction, 1.25% - 1.75% for maintenance). Core body temperature was maintained 37°C using a warm water heating system. Ventilation rate was monitored using a respiration pad (SA Instruments, USA) and maintained at 60 – 80 breaths per minute by adjustment of isoflurane concentration.

Following shimming, a T_2_-weighted, fast spin echo sequence was used for tumor localization and volume measurement (see supplemental methods for details).

#### CEST MRI

Our CEST sequence was based on a gradient echo acquisition, and included the following parameters: repetition time (TR), 162 ms; echo time (TE), 2 ms; saturation power, 3 μT; flip angle, 20°; slice thickness, 1 mm (single slice); matrix size, 64x64; FOV, 30x30 mm^2^. The slice was positioned to encompass the largest cross-sectional area of each tumor. Sampling of each line of k-space was preceded by a train of three 50 ms Gaussian pulses to induce saturation. Each pulse had a flip angle of 20°, duration of 2 ms and with 2 ms spacing (25). Crusher gradients were applied between pulses to spoil any residual transverse magnetization. The sequence was repeated with saturation frequency offsets ranging from –6 to 6 ppm, at intervals of 0.12 ppm. A reference image was also acquired at an offset of 8000 ppm (*S*_0_).

During post-processing, z-spectra were produced on a pixel-by-pixel basis, according to *Z*(*f*) = *S*(*f*) / S_0_, where *f* is the saturation offset frequency. These data were fitted with a summed Lorenzian model, with peak offsets corresponding to proton exchange between water and amide, amine and hydroxyl groups, alongside a broad peak corresponding to magnetization transfer (MT). Details of this modelling step are provided in the supplemental methods.

#### Diffusion MRI and relaxometry

Full details of diffusion MRI, T_1_ and T_2_ acquisition and quantification procedures are provided in the supplemental materials. In brief, diffusion MRI data were acquired using a multi-slice fast spin-echo sequence, with b-values ranging from 150 to 1070 s mm^2^. The apparent diffusion coefficient was quantified from these data by fitting to a simple exponential function (equ. 5). T_1_ and T_2_ were estimated from data acquired with a Look-Locker segmented inversion recovery sequence and a multi-echo multi-slice spin-echo sequence, respectively.

### *Ex vivo* analysis of resected tumor samples

Resected tumors were cut in half: one half was flash-frozen in liquid nitrogen and stored at -80 °C (for Bradford Assay) and the other half was fixed in 5% formalin for immunohistochemical analysis.

#### Bradford Assay

Protein extraction reagent was added to the tumor samples at 1g/20ml and homogenized according to the manufacturer’s instructions (T-PER^®^ Tissue Protein Extraction Reagent, Thermo Scientific). The homogenized mixture was centrifuged at 10,000 g for five minutes to pellet tissue debris. The supernatant was collected or stored at -20°C for Bradford assays. Bradford assays were carried out according to manufacturer’s instructions (Pierce Coomassie Bradford Protein Assay Kit, Thermal Scientific).

#### Immunohistochemistry

GADD34 and cleaved caspase-3 immunohistochemistry was performed on all tumor samples (details provided in supplemental methods). All slides were haematoxylin counterstained.

## Statistical Analysis

The median value for each imaging parameter was used for data analysis (rather than mean values, in order to limit the influence of extreme or outlier values resulting from convergence at local minima). Longitudinal data were normalised to the baseline measurement and expressed as a percentage change. Two-way analysis of variance between groups (ANOVA) (GraphPad Prism X.6.0.1, California, USA) was used for statistical analysis. Both the difference between control and treated groups at the same time points (between group analysis), and the difference between post-treatment and baseline data (within group analysis) were tested for statistical significance. P < 0.05 was considered significant. Correlations were assessed using Spearman’s rho.

## Results

### LS174T and SW1222 human colorectal cancer cell lines displayed differing *in vitro* sensitivities to ixazomib

The MTT assay showed that both SW1222 and LS174T cell lines demonstrate a significant decrease in viability at doses greater than 12 nM in SW1222 and 24 nM in LS174T cells (see **Figure 1**), at 24, 48 and 72 hours. Interestingly, both cell lines also exhibited a stimulatory response at low doses (<6 nM in SW1222 and <24 nM in LS174T), evidenced by an increase in cell viability (130% in SW1222 and 120% in LS174T) that peaked at 48 hours. Overall, SW1222 cells were more sensitive to ixazomib than LS174T cells (*IC*_*50*_ of 12.6 nM and 41.1 nM for SW1222 and LS174T, respectively), with an approximately three-fold lower viability at 48 hours than LS174T cells, and approximately ten-fold lower viability at 72 hours (IC_50_ of 7.6 nM and 78.4 nM for SW1222 and LS174T, respectively).

**Figure 1.**
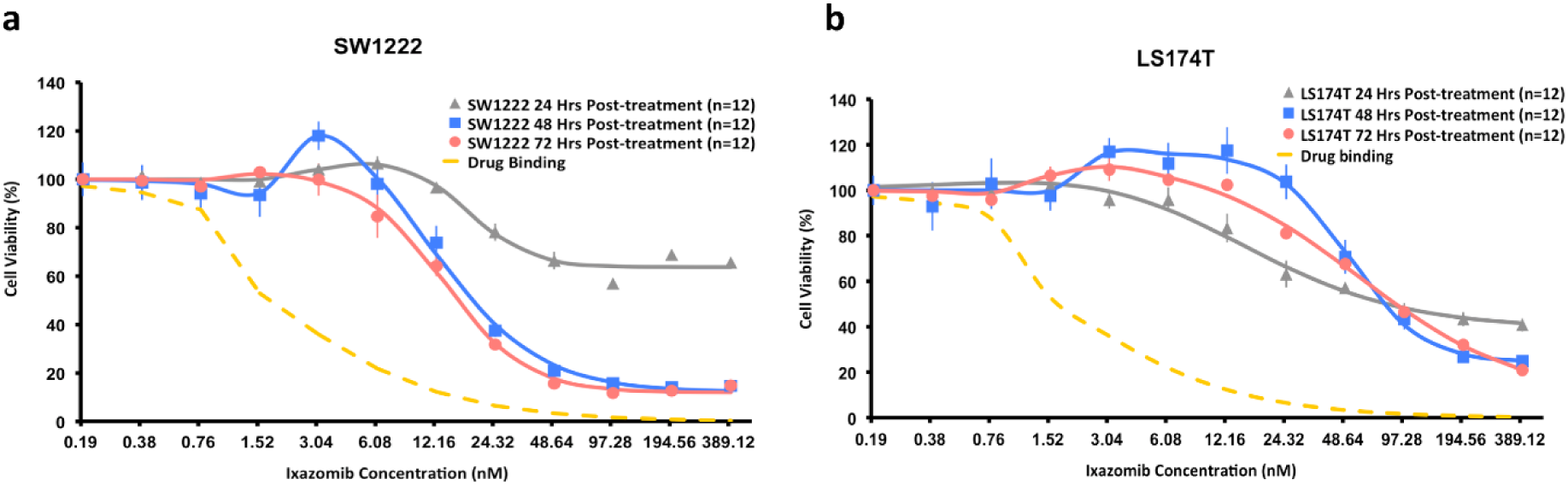
Dose-response curves showing cell viability as a function of ixazomib concentration, measured with the MTT assay, at 24, 48 and 72 hours post-dosing, in SW1222 (a) and LS174T (b) cells (data points are mean ± standard error). These *in vitro* results show that cell viability decreases at higher ixazomib doses (>12nM in SW1222 and >24 nM in LS174T) and that growth is stimulated at lower doses (<6nM in SW1222 and <24 nM in LS174T). SW1222 cells are more sensitive to ixazomib than LS174T cells, which is in keeping with their K-Ras mutation status. Dashed lines show ixazomib binding estimates. Data points are mean ± standard error.

### Ixazomib affected colorectal tumor xenograft growth *in vivo*, in a dose-dependent manner

We next investigated the action of ixazomib *in vivo*, at 24 and 72 hours after dosing with ixazomib (at 11, 9.5 or 8 mg kg^-1^) or vehicle. MRI measurements of tumor volume confirmed that the growth of both SW1222 and LS174T tumors was significantly inhibited following ixazomib doses of 9.5 and 11 mg kg^-1^ relative to control tumors (**Figs. 2** and **S1**). We also observed a small but significant increase in tumor volume at 72 hours (relative to baseline) at the lowest ixazomib dose (8 mg kg^-1^) in both SW1222 (P<0.0001) and LS174T (P<0.001) tumors, which potentially mirrors the low-dose stimulatory effect that were observed in *in vitro* experiments.

**Figure 2.**
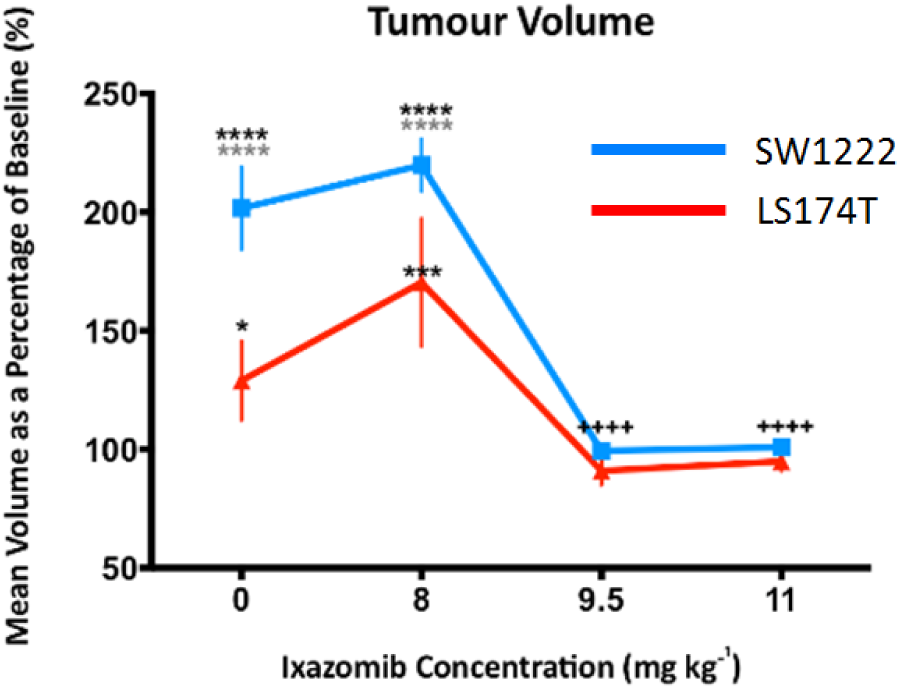
Mean relative change in tumour volume as a function of ixazomib dose, at 72 hours post-dosing, in LS174T and SW1222 human colorectal carcinoma mouse xenografts. Tumour growth was significantly inhibited at the two highest doses investigated (9.5 and 11 mg kg^-1^), and mildly (but significantly) stimulated at the lowest dose (8 mg kg^-1^). Tumour volumes were measured using volumetric MRI, and error bars represent the standard error in each measurement; * P<0.05, ** P<0.01, *** P<0.001, **** P<0.0001 (two-way ANOVA); *(black) = compared to baseline (pre-dosing) measurement, *(grey) = compared to measurement at 24 hours after ixazomib dose, ^+^ = compared to control measurement.

### CEST MRI measurements reflected response to proteasome inhibition in colorectal cancer xenografts

CEST MRI measurements were represented using Z-spectra, a plot of normalized water signal as a function of saturation frequency (**Figure 3a**). Amide, amine and hydroxyl groups are located at approximately 3.5, 2.4 and 1.2 ppm from water, respectively (25-27), alongside contributions from magnetization transfer (MT, -2.4 ppm (26)), relayed nuclear Overhouser effect (NOE, -3.3 ppm (28)) and lipids (-1 to -5 ppm (27)) combined into a single, broad peak. Fitting our summed Lorentzian model on a pixel-by-pixel basis allowed images of each z-spectrum contribution to be generated (**Figure 3b**).

**Figure 3.**
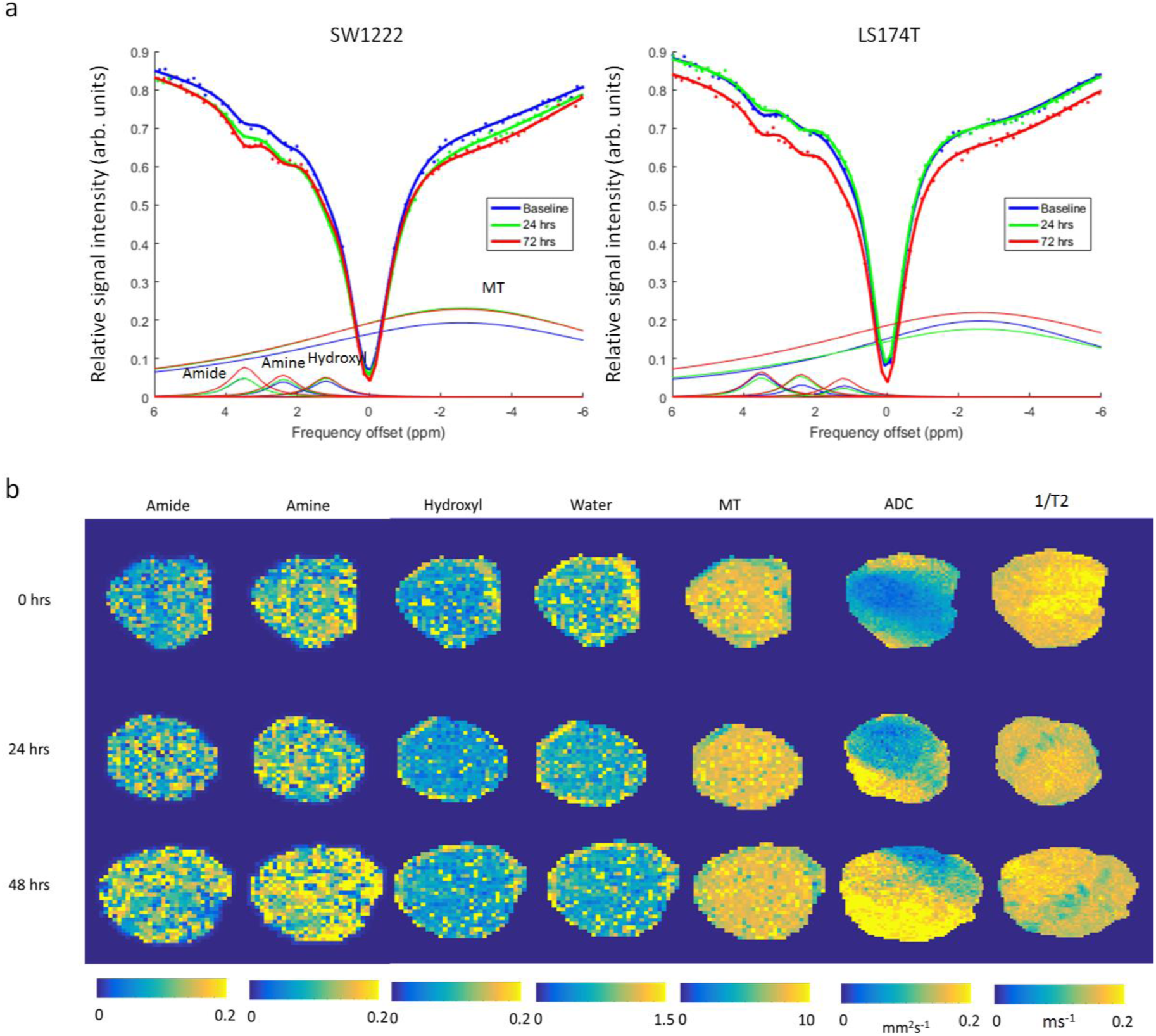
Example MRI measurements in SW1222 and LS174T tumours, prior to and at 24 and 72 hours following treatment with ixazomib, showing increasing amide and amine peak areas, alongside increasing apparent diffusion coefficient (ADC). (a) Z-spectra from CEST MRI, showing the normalised MRI signal intensity as a function of saturation frequency, averaged over a single representative tumour dataset at baseline and 24 and 72 hours following dosing with 11 mg kg^-1^ of ixazomib. Data points were fitted to a Bayesian model that allowed the individual influence of proton exchange between water and amide, amine and hydroxyl groups to be isolated (1.2, 2.4 and 3.5 ppm, respectively), alongside magnetisation transfer (MT, -2.4 ppm). Water peaks (0 ppm) have been removed for clarity. (b) Images of amide, amine hydroxyl, MT and water peak areas, acquired noninvasively in an example SW1222 tumour at baseline and post-therapy (24 and 72 hours). Maps of the apparent diffusion coefficient (ADC) from diffusion MRI measurements and the transverse relaxation time (T_2_) are also shown.

The area under amide and amine peaks significantly increased following ixazomib treatment, relative to pretreatment measurements (**Figs. 4** and **S1**). A summary of all significant changes, for each tumor type, ixazomib dose and time post-dosing is provided in **Table 2**. These data, in combination, demonstrate the smaller magnitude of change in parameter values following treatment in LS174T tumors compared with SW1222 tumors, potentially reflecting their lower sensitivity to ixazomib.

**Figure 4.**
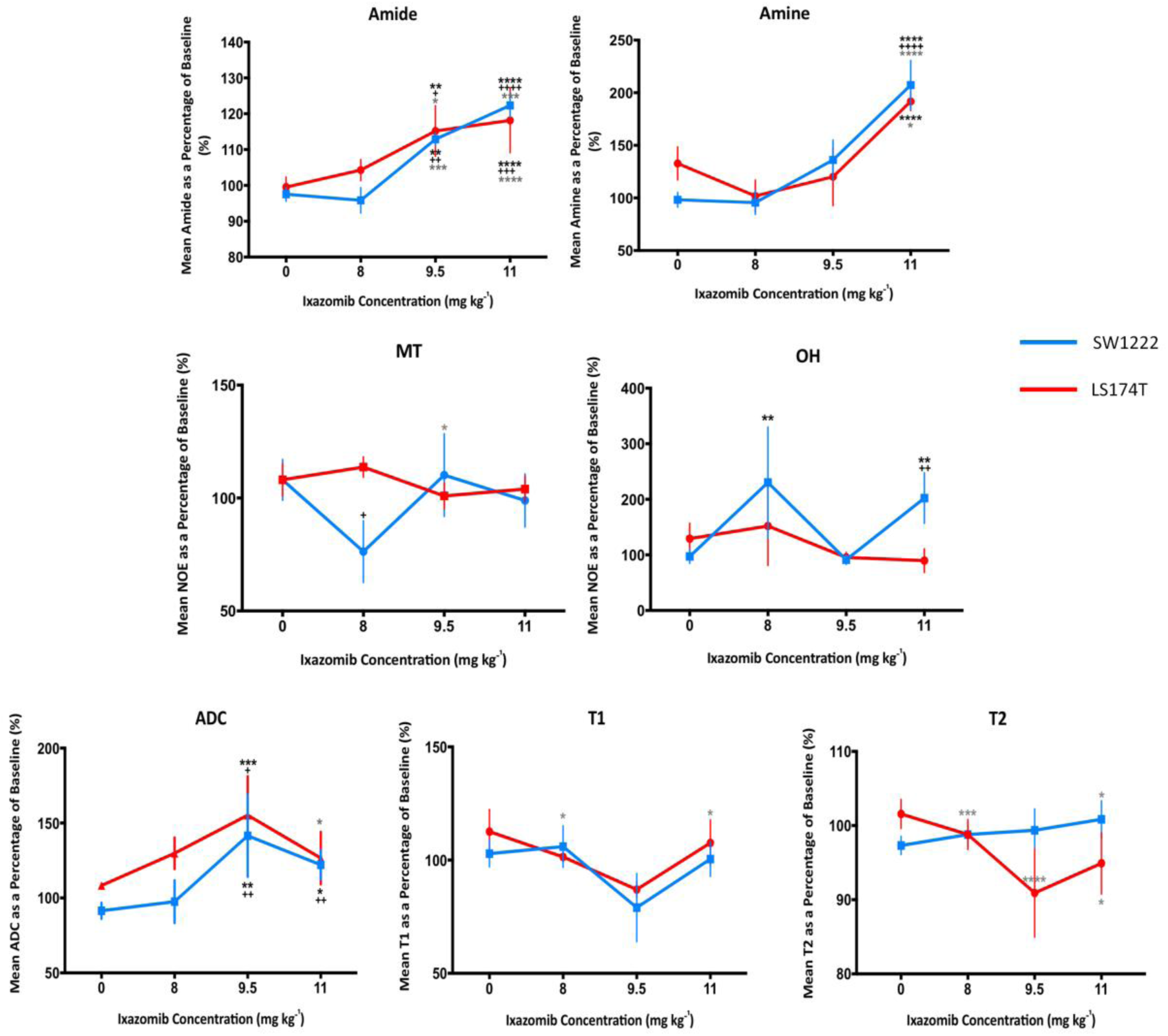
Summary of changes in quantitative, non-invasive MRI parameters, as a function of ixazomib dose (8, 9.5 and 11 mg kg^-1^), at 72 hours post-dosing. Only CEST measurements associated with amines and amide groups show a correspondence with ixazomib dose. Significant increases, in a dose-dependent manner, were observed in amine and amide peak areas from CEST MRI measurements. Tumour apparent diffusion coefficient (ADC) also increased significantly for the two highest doses, and T_2_ decreased significantly in LS174T tumours only. Data points are mean ± standard error; * P<0.05, ** P<0.01, *** P<0.001, **** P<0.0001 (two-way ANOVA); *(black) = compared to baseline, *(grey) = compared to 24 hours post-ixazomib dose, ^+^ = compared to control.

### Changes in amide and amine peaks were significantly correlated with ixazomib dose

A significant correlation was found between amide peak area and ixazomib dose in SW1222 tumors at 72 hours (r = 0.98, P< 0.05) (**Figure 4**). This was also reflected in the change in the amine area of the Z spectrum, with a strong positive correlation between the dose concentration and amine signals (r=0.97, P<0.05). Conversely, no parameters in LS174T tumors exhibited a dose-dependent response, which could reflect their reduced sensitivity to ixazomib, compared with SW1222 tumors.

### Gross tissue protein measurements from Bradford assay displayed a complex relationship with ixazomib dose

Bradford assay at 72 hours showed a significant difference in mean protein concentration between SW1222 and LS174T control tumors (1386 μg ml^-1^, P<0.001). LS174T tumors had a small but significantly lower protein concentration than control tumors when dosed at 8 mg kg^-1^, whereas higher doses did not induce a significant change (**Figure 5c**). SW1222 tumors exhibited lower gross protein concentration when treated at 8 and 11 mg kg^-1^, compared with control tumors, but a significantly higher protein concentration at 9.5 mg kg^-1^.

**Figure 5.**
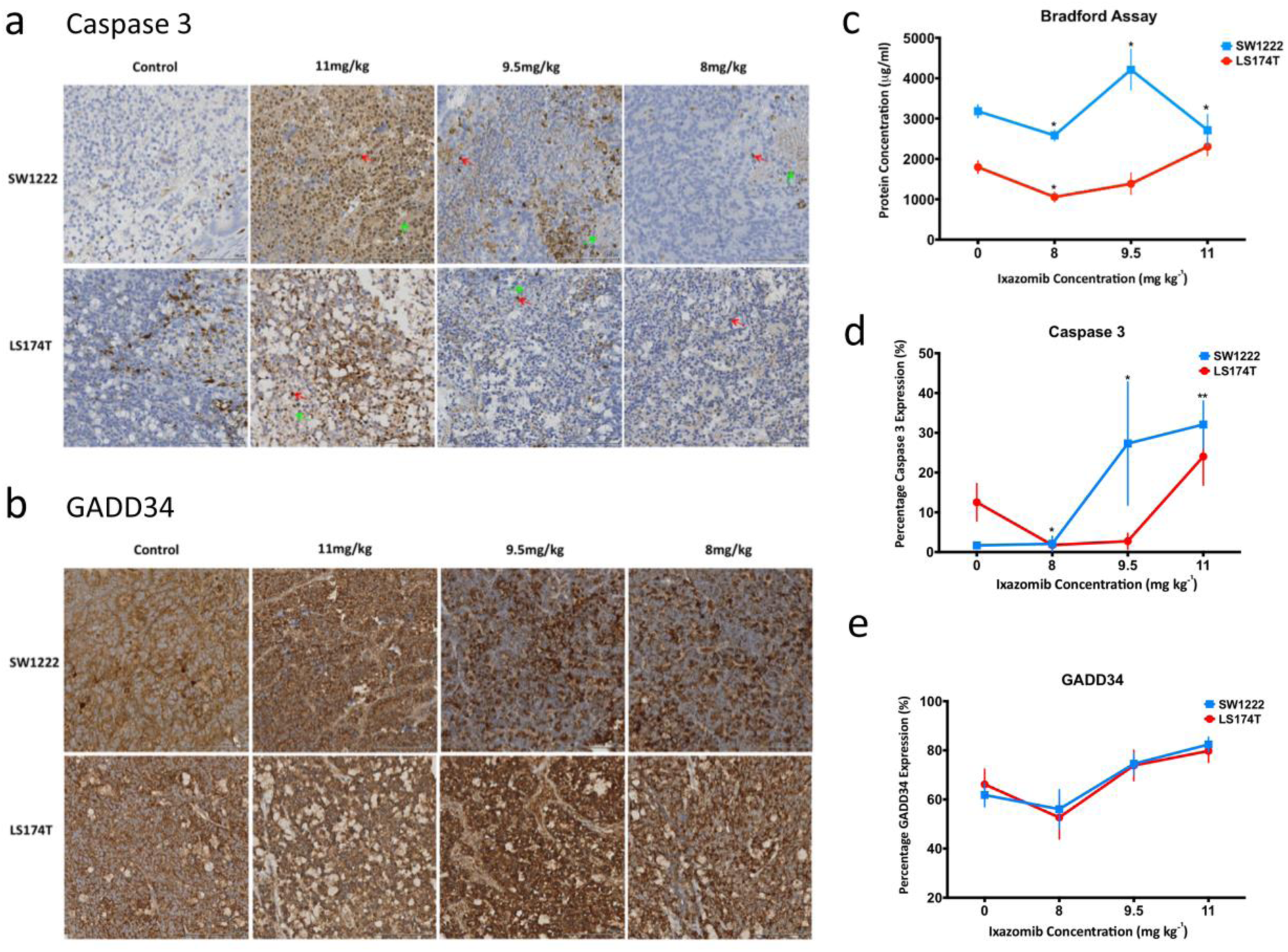
*Ex vivo* analysis of resected tumour tissue. Images of (a) caspase 3 and (b) GADD34 expression at 72 hours following dosing with vehicle, 8mg kg^-1^, 9.5mg kg^-1^ or 11 mg kg^-1^. Caspase 3-positive nuclei are stained brown (as labelled with a red arrow) and blue otherwise (haematoxylin), and example necrotic areas are labelled with a green arrow. GADD34-positive areas are stained brown and nuclei are counter-stained in blue with haematoxylin. (c) The results of Bradford assay, to measure gross tissue protein. Summary data from quantitative analysis of immunohistochemistry images are shown for caspase 3 (d) and GADD34 (e), for each study group. Data points are mean ± standard error (* P<0.05, ** P<0.01, ordinary one-way ANOVA of Mann- Whitney test).

No significant correlation was found between Bradford assay results and *in vivo* amide and amine peak area (P>0.05, Spearman’s rho). Equally, no significant differences were found in baseline CEST parameters, between tumors, that might have reflected the difference found between tumor types in Bradford assay results.

### Diffusion MRI measures reflected changes in CEST parameters, but T1 and T2 showed mixed changes

A significant increase was found in ADC from baseline, in SW1222 tumors at 72 hours, for ixazomib doses of 9.5 and 11 mg kg^-1^ (41.5%, P<0.01 and 22.1%, P<0.05, respectively), which were significantly greater than control tumors (**Figure 4** and **Table 2**). The change in ADC from baseline was also significantly correlated with ixazomib dose (P<0.01). In LS174T tumors, only the 9.5 mg kg^-1^ group showed a significant increase, when compared against baseline (55.19%, P<0.001) and control tumors (46.92%, P<0.05).

T_1_ and T_2_ showed a more mixed set of changes in both tumor types, with fewer time points achieving significance (see **Figure 4** and **Table 2**). T_1_ and T_2_ showed both increases and decreases over time, and did not display a simple relationship with ixazomib dose, other than for the change in T_2_ in SW1222 tumors at 72 hours, which was directly correlated with dose (r = 0.99, P<0.01).

**Table 1.** Gaussian prior distribution parameter values for the maximum likelihood model, expressed as mean (s.d.) (derived from Chappell *et al*. (26)). Entries marked with ‘n/a’ were fixed (see text for details).

| Pool (index) | Gaussian prior parameters: mean (s.d.) |  |  |
| --- | --- | --- | --- |
|  | *b* (ms) | *a* (ppm) | *f<sub>o</sub>* (ppm) |
| Water (0) | 5 (3) | n/a | 0 (3) |
| Amide (1) | 8 (5) | 3.5 (0.6) |  |
| Amine (2) | 8 (5) | 2.4 (0.6) |  |
| Hydroxyl (3) | 5 (5) | 1.2 (0.6) |  |
| MT & NOE (4) | 0.1 (1) | -2.4 (0.6) |  |

**Table 2.**
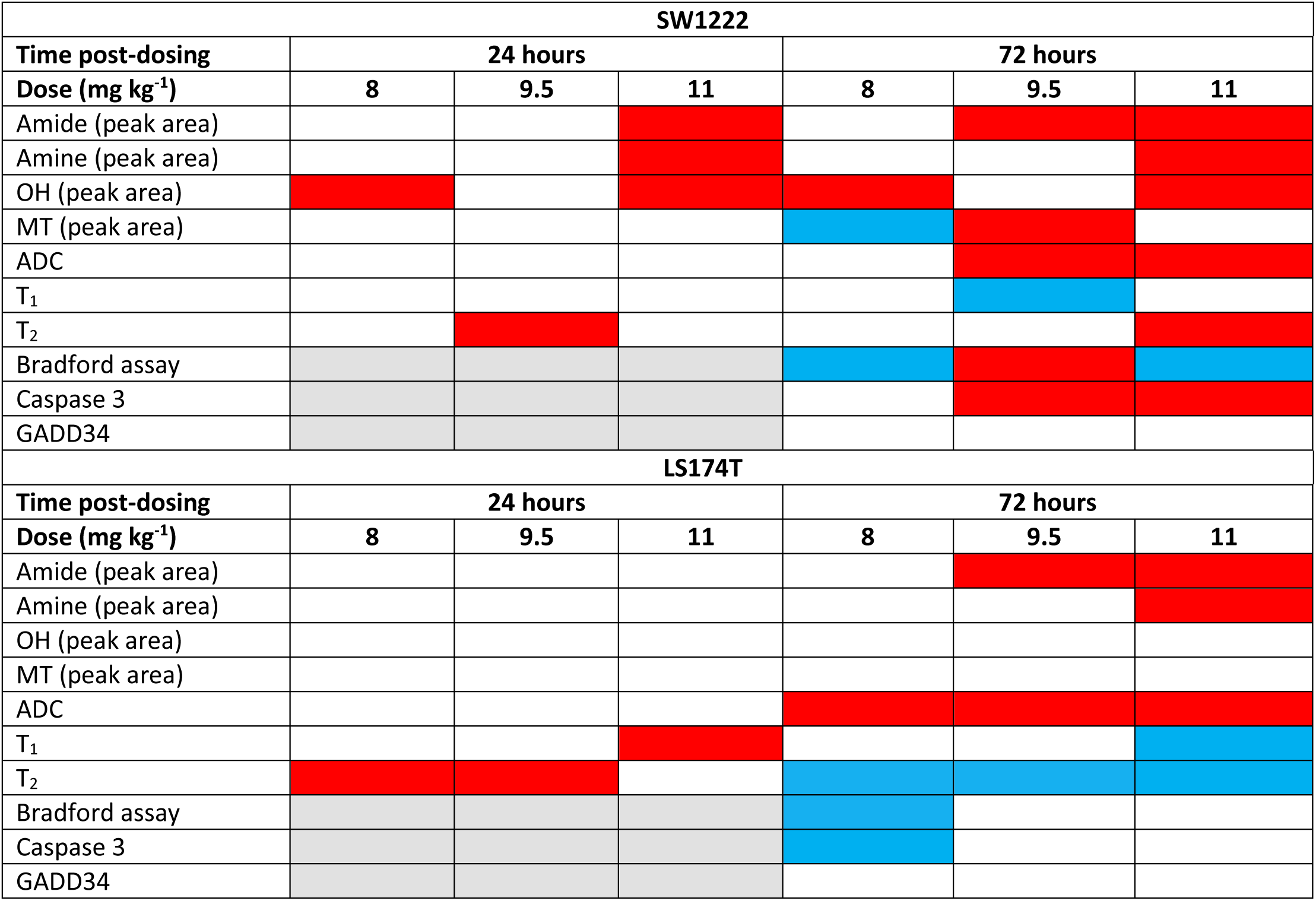
Summary of significant changes in each quantitative parameter measured, relative to controls. Red cells indicate a significant increase, blue cells a significant decrease, white cells no significant change, and grey cells show where no data were available.

### Immunohistochemical markers of apoptosis significantly correlated with ixazomib dose in SW1222, but not LS174T tumors

To evaluate the relationship between CEST imaging biomarkers and physiological changes caused by proteasome inhibition, we used immunohistochemistry to measure the percentage expression of cell death factor, caspase 3, and GADD34 (20, 29). Our results revealed significantly higher caspase 3 expression in SW1222 tumors for doses of 9.5 mg kg^-1^ (25.61%, P<0.05) and 11mg kg^-1^ (29.87%, P = 0.002), relative to controls (**Figure 5**). Caspase 3 expression was also significantly correlated with ixazomib dose (r=0.95, P<0.05). Though small, there was significantly lower caspase 3 expression in LS174T tumors for a dose of 8 mg kg^-1^ (4.57%, P = 0.0127), relative to controls, but no significant differences at higher doses. Whilst there was no significant change in GADD34 percentage expression relative to controls in either SW1222 or LS174T tumors (**Figure 5**), as for caspase 3, a significant correlation was found between GADD34 expression with increasing dosing concentration in SW1222 tumors (r = 0.96, P<0.05).

## Discussion

Noninvasive imaging biomarkers for cancer therapy are critically needed in the clinic, particularly those that can detect a functional response prior to much slower changes in tumor volume (22). CEST is a noninvasive imaging technique, based on MRI, in which image contrast is induced via the exchange of protons between water and chemical groups such as amides, amines and hydroxyls (17). The presence of amide groups in protein backbones and amine groups on peptides make CEST sensitive to changes in the concentration of both types of molecules (30-34), and therefore as a potential biomarker of disrupted protein homeostasis. We therefore aimed to evaluate the change in CEST MRI parameters (alongside other more conventional MRI measures) following dosing with ixazomib, in two colorectal carcinoma tumor xenograft models (SW1222 and LS174T).

Our experiments showed that ixazomib elicited a significant growth-inhibitory effect in both colorectal cell lines, as evidenced by a reduction in cell viability *in vitro* and in the reduction of tumor growth rates *in vivo*. SW1222 tumors were found to be more sensitive to ixazomib than LS174T tumors, which could be due to their different K-Ras mutations: SW1222 cells have a K-Ras mutation at A146V (35), compared with G12D for LS174T tumors (36), which has been proposed to infer a greater sensitivity to proteosome inhibition (11). In addition, the microenvironment of the two tumor types could provide a source of resistance *in vivo*, with LS174T tumors being less vascular, less differentiated and less perfused than SW1222 tumors (37-39).

Our CEST measurements showed a significant increase in amide, amine and OH peak areas with time following ixazomib treatment, with the greatest effects observed at 72 hours following dosing. Moreover, several parameters varied in a dose-dependent manner, with a significant correlation measured between amide and amine peak size and ixazomib dose in SW1222 tumors. This reflected similar trends in our immunohistochemical measurements of caspase 3 and GADD34 expression, which were themselves consistent with previous studies (20, 29). This induction of apoptosis could also explain the observed increases in ADC, in which changes in cell size and the induction of micro-necrosis causes less restricted water diffusion (37).

However, the relationship between CEST parameters and Bradford assay were less straightforward to interpret, with no apparent correspondence evident between the two measurement types. Several factors might help to explain this disparity. Firstly, CEST MRI is sensitive to mobile protein and peptides, whereas the Bradford assay is unaffected by amino acids or peptides smaller than 3 kDa (according to manufacturer’s instructions). Secondly, CEST contrast is dependent on several other parameters including *T*_1_, *T*_2_ and pH, which were not controlled for in this study. *T*_1_ and *T*_2_ were found to show few consistent changes when measured directly, so it is arguable that these were unlikely to have influenced CEST measurements significantly. The pH of tumor tissue could also have influenced our measurements, and a fully quantitative model (26) could potentially be implemented to separate proton exchange rate and pool sizes.

Interestingly, we found evidence that ixazomib causes a mild stimulatory effect at low doses (40). This bi-phasic effect has been previously observed in a wide range of anticancer agents (40-43) alongside physical interventions such as radiotherapy (44). *In vitro* MTT assays revealed increased cell viability at low doses, and tumor growth was slightly enhanced at an *in vivo* dose of 8 mg kg^-1^. Likewise, caspase 3 expression was slightly decreased, relative to controls, in LS174T tumors dosed at 8 mg kg^-1^. Whether this effect was reflected in MRI measurements is unclear, as no significant changes were observed in CEST parameters (amide and amine areas) at low ixazomib doses. However, this effect warrants further investigation, particularly if the treatment of tumors with poor delivery profiles (for example, pancreatic adenocarcinoma) is currently under consideration, as delivered doses might be significantly lower than expected. In this context, in particular, amide and amine CEST signals and ADC could find strong utility, alongside complementary measures of blood flow and interstitial transport.

A key question is whether these findings can be translated and reproduced in humans. Clinical MRI is typically undertaken at a field strength of 3T, which is lower than the 9.4T system used here. Translation between field strengths presents several challenges (such as reduced separation between exchange peaks, lower signal-to-noise ratios, and slower relaxation rates), but which have been overcome in other translational studies. Equally, the evaluation of Ixazomib in human solid tumors is at an early stage and the data presented here demonstrates its efficacy in two colorectal carcinoma xenograft models. It also identifies several promising, noninvasive biomarkers (amide and amine CEST signals and ADC) that could be explored for determining early treatment response and detecting disrupted protein homeostasis. In addition, they could be further evaluated in the assessment of plasma cell malignancies (currently the predominant targets of Ixazomib), with a view to reducing the need for invasive bone marrow aspirates or biopsy.

## Supplemental methods

### Cell culture

SW1222 and LS174T cells were cultured in minimum essential medium eagle (MEME, M4655, Sigma-Aldrich Cell Culture, Gillingham, UK) with 5% fetal bovine serum. On the day of inoculation, the cells were harvested by washing once in phosphate buffered saline (PBS) solution (Sigma-Aldrich Cell Culture, Gillingham, UK) and trypsinised at 37°C for 5 minutes. The cells were pelleted by suspending in MEME and centrifuged at 1600 x g for 5 minutes. The suspension mixture was removed and the pelleted cells were washed in PBS and then in serum-free media at 1600 x g for 5 minutes. Before the final wash in serum-free medium, 10 μl of the cell suspension mixed with trypan blue (Invitrogen) was transferred to a cell counting chamber slide (Invitrogen) and cells were counted with a Countess Automated Cell Counter (Invitrogen). The cells pelleted after the final wash were suspended in known volume serum free-media.

### MTT assay

#### Quantification

We estimated cell viability from the absorbance measure (unitless), in which the viability of treated cells was expressed as a percentage of the mean absorbance of control cells. Mean measurements of cell viability, expressed as a percentage change from baseline, were fitted to the modified Hill equation:

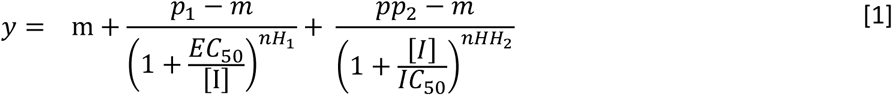

where *m* is the maximum percentage viability; *p1* is the viability at low concentration (constrained to 100%), and *p*_*2*_ is the viability at higher concentrations; [*I*] is the concentration of ixazomib (nM); *EC*_*50*_ and *IC50* are concentrations that produced half-maximal stimulatory and inhibitory effects (nM), respectively; *nH*_*1*_ *and nH*_*2*_ are unitless Hill factors for stimulatory and inhibitory slopes, respectively.

The modified Hill equation describes the binding of a drug to a receptor, with an empirical modification factor (*nH*) (1). In our implementation, we modeled a single ixazomib binding site on the proteasome. The relationship between binding the proteasome and inhibition of cell proliferation is unknown. The data were fitted with a two-component Hill Equation to estimate the *IC*_*50*_ concentration and maximum effect. Data were fitted using Prism (GraphPad Prism X.6.0.1, California, USA).

### CEST z-spectrum model

The Lorentzian function for the *i*^*th*^ pool was given by:

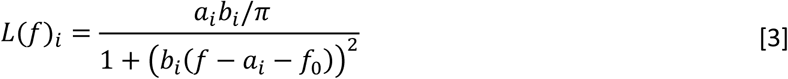

which were fitted to z-spectra according to:

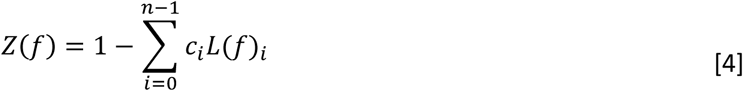

*a*_*i*_ is the estimated frequency offset of the *i*^*th*^ pool, *bi* is the width parameter, *c*_*i*_ is a pool-specific scaling factor and *f*_0_ is a global frequency offset to account for inhomogeneities in the static magnetic field (*B*_0_). Bayesian maximum likelihood was used to fit *a, b c* and *f*_0_ to z-spectra (2), using the fminunc minimization algorithm in Matlab (MathWorks, Massachusetts, US) with the exception of a_0_ (water pool), which was fixed at 0 ppm. Gaussian prior distributions for each parameter were defined as shown in **Table S1,** which were derived from Chappell *et al*. (3).

### Tumor volume measurement with MRI

A T_2_-weighted, fast spin echo sequence was used for tumor localization and tumor volume measurements, which included the following parameters: repetition time (TR), 1.5s; echo train length, 4; effective echo time (TE_eff_), 17.2 ms; slice thickness, 1mm; number of slices, 20; number of averages, 4; matrix size, 128x128; field of view (FOV), 30x30 mm^2^. Regions of interest (ROIs) were drawn, covering the entire tumor mass (Matlab). The number of pixels in these volumetric ROIs was multiplied by the voxel volume (238×238×1000 μm^3^) to estimate tumor volume.

### Diffusion MRI acquisition and post-processing

A multi-slice diffusion-weighted fast spin echo sequence was used which included the following parameters: TR, 1500 ms; TE_eff_, 2000 ms; b-value, 150, 300, 503, 760, 1070 s mm^-2^; slice thickness, 1 mm; number of slices, 8; matrix size, 128x128; FOV, 30x30 mm^2^. The apparent diffusion coefficient (ADC) was quantified from these data by fitting a single exponential to signal intensity values, of the form:

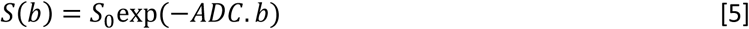

where *S*_0_ and ADC are fitted parameters and *b* is the b-value. This was undertaken on a pixel-by-pixel basis, and pixels were fitted using a maximum likelihood algorithm that took into account the Rician noise distribution (4).

### *T*_1_ measurement

A Look-Locker segmented inversion recovery sequence was used to estimate the longitudinal relaxation time, *T*_1_, with the following parameters: number of inversion points, 50; TR, 110 ms; TE, 1.18 ms; slice thickness, 1mm; number of slices, 8; matrix size, 128x128; FOV, 30x30 mm^2^. In order to sample the longitudinal magnetization during its recovery following an inversion pulse, the Look-Locker sequence applies a train of small flip angle readout pulses, separated by a fixed inversion time (TI). This pulse train partially saturates the signal, resulting in an apparent longitudinal relaxation time, *T*_1_*, shorter than the true *T*_1_ time. *T*_1_* was estimated by fitting data to the following three parameter model:

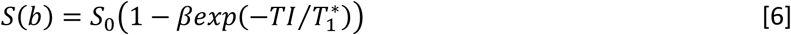

*T*_1_ was then estimated according to (5)

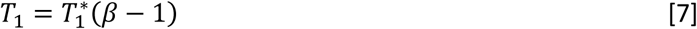

### *T*_2_ measurement

A multi-echo multi-slice imaging sequence was used to estimate *T*_2_, using the following parameters: TR, 1500 ms; 16 echo times (TE) ranging from 8 to 132 ms, at 8 ms intervals; slice thickness, 1 mm; number of slices, 8; matrix size, 128x128; FOV, 30x30 mm^2^. *T*_2_ and *S*_0_ values were estimated using the following equation:

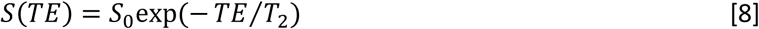

where S_0_ and *T*_2_ are fitted parameters. Again, a maximum likelihood algorithm was used, which took into account the Rician noise distribution of MRI magnitude data (2).

### Immunohistochemistry

#### Staining and Imaging

GADD34 and cleaved caspase-3 immunohistochemistry was performed using a Ventana Discovery XT, with the Ventana DAB Map Kit (760-124). Details of the protocols used are provided in supplemental methods. Heat induced epitope retrieval was performed using an EDTA buffer (pH8.0). Anti- GADD34 (Santa Cruz Biotech SC-8327) primary antibody incubation was for 4 hours using a 1:50 dilution, followed by amplification with the Ventana Amplification Kit (760-080). Swine anti-Rabbit (Dako E0353) secondary antibody incubation was for 30min, using a 1:200 dilution. Anti-Cleaved-Caspase-3 (Cell Signaling, 9661L) primary antibody incubation was for 30 minutes using a 1:100 dilution. Swine anti-Rabbit (Dako E0353) secondary antibody incubation was for 30 minutes, using a 1:200 dilution. All slides were haematoxylin counterstained and digitalized using LEICA SCN400 scanner (LEICA Microsystems, Milton Keynes, UK), with x40 magnification, 65% image compression.

#### Analysis and quantification

Slides were analyzed and quantified using Definines Tissue Studio and Developer (Definines AG, Munich, Germany). Marker expression was quantified by measuring percentage area and percentage nuclei of the positive regions. For the percentage area calculation, distinctive regions of a tumor tissue slide were first manually classified from a sample slide. A typical immunostained tumor tissue was classified into five regions: positive, viable, necrotic, dermis/fat, and glass, in the training process of Tissue Studio composer actions. The software then automatically identified and segmented each slide into five regions of interest (ROI) with threshold based on the intensity of haematoxylin (blue) and chromogen (brown). After the automatic segmentation of all slides was completed, the slides were reassessed manually by the user to correct any mismatched regions (in ‘correction mode’). Automatic quantification was then carried out, where the absolute area and percentage area of each ROI over total tissue was calculated, where the total tissue area was the sum of positive, viable and necrotic areas. For the percentage nuclei quantification, in the software training process, tumor tissue area (from segmentation described above) were classified as ‘nuclei’ or ‘cytoplasm’, and within the ‘nuclei’, they were further classified into ‘positive’, ‘viable’ and ‘necrotic’. The tissues were automatically labeled based on the preset threshold values of the stain intensity. The absolute number and percentage of each class of nuclei were automatically quantified, where the total of nuclei were the sum of positive, viable and necrotic nuclei. The percentage expression of caspase 3 and GADD34 were calculated by taking a mean value between the percentage area and percentage nuclei.

---

**Table S1.** Gaussian prior distribution parameter values for the maximum likelihood model, expressed as mean (s.d.) (derived from Chappell *et al*. (26)). Entries marked with ‘n/a’ were fixed (see text for details).

| Pool (index) | Gaussian prior parameters: mean (s.d.) |  |  |
| --- | --- | --- | --- |
|  | *b* (ms) | *a* (ppm) | *f<sub>0</sub>* (ppm) |
| Water (0) | 5 (3) | n/a | 0 (3) |
| Amide (1) | 8 (5) | 3.5 (0.6) |  |
| Amine (2) | 8 (5) | 2.4 (0.6) |  |
| Hydroxyl (3) | 5 (5) | 1.2 (0.6) |  |
| MT & NOE (4) | 0.1 (1) | -2.4 (0.6) |  |

**Figure S1.**
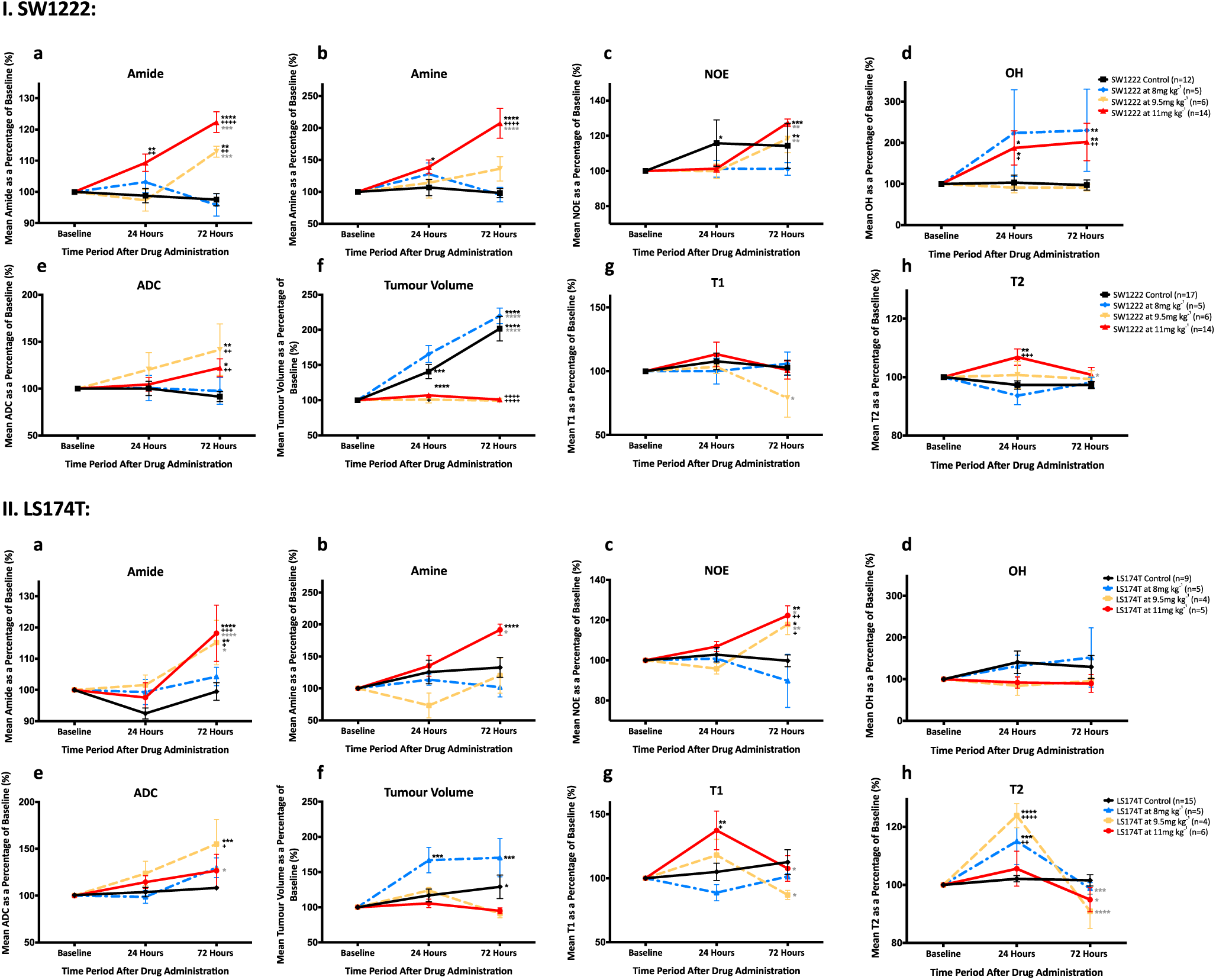
Graphs showing the relationship between the relative change in each non-invasive imaging parameter and time, for three different doses of Ixazomib (8, 9.5 and 11 mg kg^-1^) and vehicle, in (I) SW1222 and (II) LS174T tumours. Imaging parameters shown are: (a) median amide area, (b) median amine area, (c) median NOE area, (d) median OH area, (e) median ADC, (f) median volume, (g) median T1 and (h) median T2. Error bars correspond to the standard error of the mean (*P<0.05, *** P<0.01, **** P<0.001; *(black) = compared to baseline, *(grey) = compared 24 hours after ixazomib treatment, ^+^ = compared to control).

